# A supergene in seaweed flies modulates male traits and female perception

**DOI:** 10.1101/2021.06.30.450538

**Authors:** Swantje Enge, Claire Mérot, Raimondas Mozūraitis, Violeta Apšegaitė, Louis Bernatchez, Gerrit A. Martens, Sandra Radžiutė, Henrik Pavia, Emma L. Berdan

## Abstract

Supergenes, tightly linked allelic combinations that underlie complex adaptive phenotypes represent a critical mechanism protecting intra-specific polymorphism [1, 2]. Supergenes represent some of the best examples of balancing selection in nature and there is increasing evidence that disassortative mating, when individuals preferentially mate with dissimilar phenotypes, is a key force stabilizing supergene polymorphisms. Yet, the underlying biological mechanisms and genetic basis of disassortative mating remain poorly known. Here, we examine a possible mechanism of disassortative mating driven by female mate choice in relation to the overdominant *Cf-Inv(1)* supergene in the seaweed fly *Coelopa frigida* by investigating chemical communication and its genomic architecture. We show that *Cf-Inv(1)* strongly affects chemical signaling; cuticular hydrocarbon (CHC) composition differed between genotypes in males but not females across two continents. In tandem, *Cf-Inv(1)* affected female perception of these compounds; females are able to sense 36 compounds from the male CHC cocktail but show differential perception between genotypes for almost half of them. This indicates that the genetic underpinnings of male traits and female perceptions are tightly linked within *Cf-Inv(1)* which likely facilitates disassortative mating [3]. A differential expression approach based on candidate genes for CHC biosynthesis and odorant detection revealed differential expression for CHC biosynthesis in males alone but broad changes in odorant receptors across both sexes. Furthermore, odorant genes clustered together within *Cf-Inv(1)*, with some of them differing between arrangements by 8.3% at the protein level, suggesting evolution via tandem duplication then divergence. We propose that the tight linkage between overdominant loci, male traits, and female perception has helped to maintain the *Cf-Inv(1)* polymorphism across its range in the face of supergene degeneration.

## Results and Discussion

Complex multi-trait polymorphisms are a fascinating aspect of intra-specific diversity. Such polymorphisms are increasingly known to be associated with supergenes, *i.e*. genomic regions harboring linked combinations of alleles from recombination [2, 4, 5]. Yet the persistence of supergene polymorphism over long time scales is puzzling especially given that supergene haplotypes often degrade [e.g., accumulate deleterious mutations; 6, 7–9]. While several mechanisms of balancing selection have been identified in classic supergene polymorphisms one form, disassortative mating (when individuals preferentially mate with dissimilar phenotypes) appears to be quite prevalent [7]. Disassortative mating, occurs in many classic supergene systems [10–14] and is adaptive in degraded supergenes because deleterious mutations are generally private to their supergene haplotype, generating a heterozygote advantage [3, 6, 15, 16]. The evolution of such a mating system requires either (i) a self-referencing system where individuals use their own phenotype to choose a mate or (ii) tight linkage between mating signals and preferences [3, 17]. By including many loci, supergenes are obviously good candidates for the latter situation, yet the modalities and the genetic architecture of disassortative mating in supergenes remain largely unknown.

To better understand this form of mate choice and its role in supergene maintenance, we explored the traits and genetic basis of disassortative mating in relation to the *Cf-Inv(1)* supergene in the seaweed fly *Coelopa frigida. Cf-Inv(1)* is a large supergene spanning 60% of chromosome 1 and 10% of the genome [18] with two arrangements, termed α and β, resulting from 3 overlapping inversions [19]. While mating appears disassortative with respect to *Cf-Inv(1)* [20–22], males perform no courtship displays in this system. Instead, mating in *C. frigida* is characterized by a sexual struggle in which females engage in various behaviors to dislodge mounted males [20, 21, 23], and experimental evidence suggests active female choice [20, 21]. Thus, disassortative mating in *C. frigida* is likely a function of female choice as males appear to mount females indiscriminately and there is strong evidence for convenience polyandry in this system [24]. As females receive no visual cues, we hypothesized that chemical cues may play a role in sexual communication in this system. Cuticular hydrocarbons (CHCs) are a common mode of communication in insects and often facilitate mate choice [25, 26], and our previous work has demonstrated its importance in seaweed flies [27]. Therefore, we investigated differences in CHC profiles between *Cf-Inv(1)* genotypes and predicted that male CHC profiles might show a stronger effect of supergene genotype than female profiles.

To quantify the role of *Cf-Inv(1)* in CHC composition, we collected CHC profiles using gas chromatography and mass spectrometry (GC-MS) from a total of 276 *Coelopa frigida* and analyzed variation between sexes and genotypes. To ask whether putative differences are conserved, we included populations from both North America (Kamouraska, QC, Canada 47.56294, −69.87375) and Europe (Østhassel, Norway 58.07068, 6.64346). The samples included 20-25 individuals of each sex, genotype (αα, αβ, ββ), and population combination (see Table S1 for a full summary of sample sizes).

The overall CHC composition of males, analyzed by PCA, shows robust differentiation between the αα and ββ genotypes on both continents, with the αβ genotype as an intermediate phenotype (Fig. S1). The effect of genotype (αα vs. ββ) on CHC composition was further investigated by OPLS-DA and PERMANOVA (Table S2). OPLS-DA separates variation in CHC composition into variation correlated with genotype (the predictive component = between group variation) and other systemic variation uncorrelated with genotype (the orthogonal components = within group variation). In males, the two homozygous genotypes showed parallel separation by the OPLS-DA models on both continents (Norway R2Y=0.91, Canada R2Y=0.92, Fig. 1A), which also showed reliable predictive performances (Norway Q2=0.57, Canada Q2=0.83, Fig. 1A). Furthermore, 19% (Canadian males) and 24% (Norwegian males) of the total variation in CHC composition contributed to the separation of the αα and ββ genotypes. The effect of the supergene on CHC composition was further supported by the PERMANOVA results (Fig. 1B) demonstrating a significant, parallel difference (α=0.05) between *Cf-Inv(1)* genotypes in both Norwegian and Canadian males.

**Figure 1 -.**
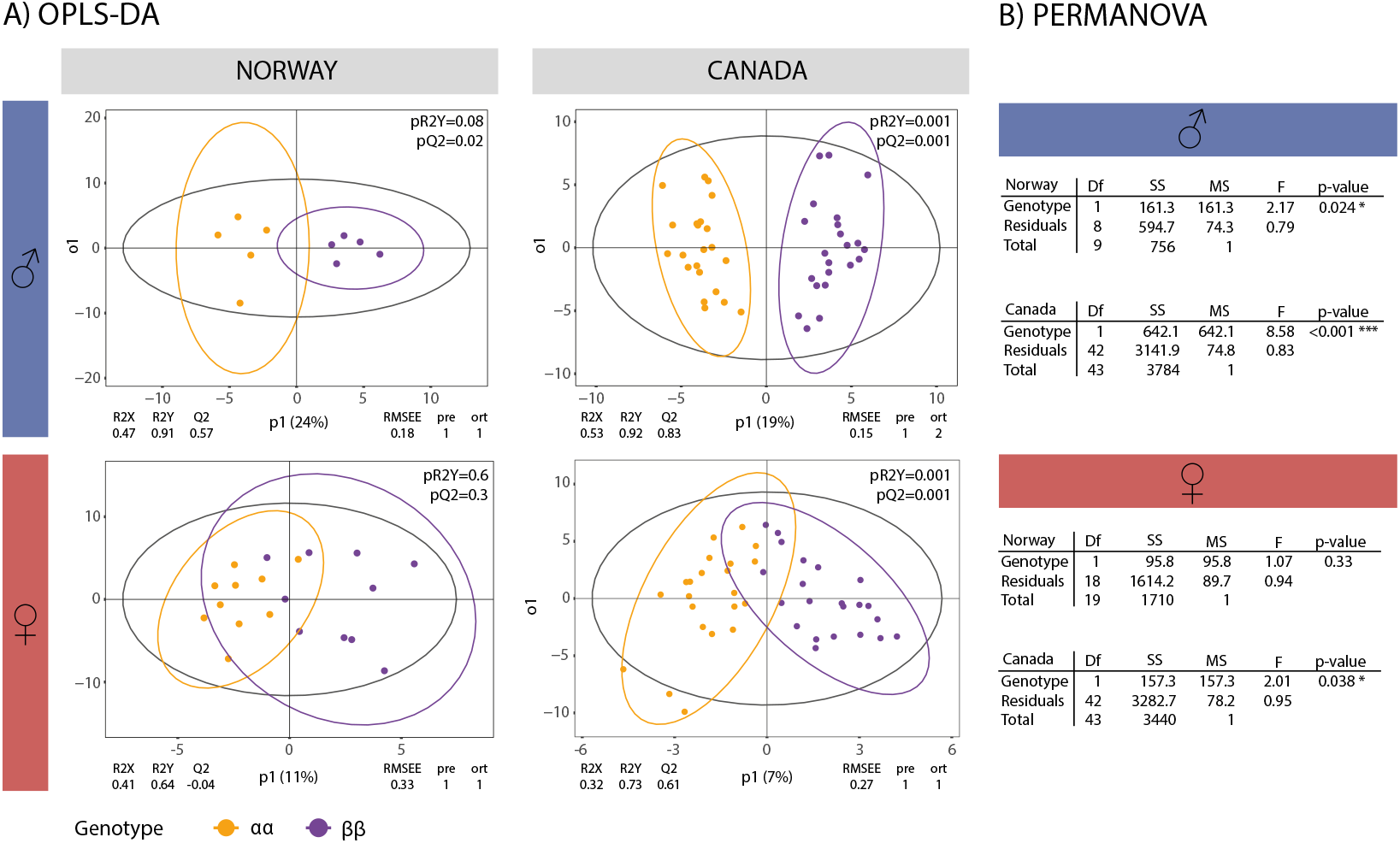
CHC composition varies by genotype in males but not females. OPLS-DA (A) and PERMANOVA (B) analysis of CHC composition. (A) OPLS-DA analysis of CHC composition in Norwegian and Canadian populations. Figures are divided by population and sex and colored by genotype: orange - αα, grey - αβ, purple - ββ. (B) PERMANOVA results. Models were run separately for each population × sex combination. Only αα and ββ individuals were used in all analyses.

In contrast, very little variation in female CHC composition could be explained by *Cf-Inv(1)* genotype. The overall CHC composition of females, analyzed by PCA, showed no differentiation between genotypes in both populations (Fig. S1). The OPLS-DA approach revealed that only 7% of the total variation in CHCs was associated with the genotype in the Canadian females. Despite reliable predictive performance (Q2=0.61), the separation of the genotypes was less pronounced than in males (RY2=0.73, Fig. 1A). In the Norwegian females, no reliable predictive OPLS-DA model was obtained (RY2=0.64, Q2=-0.04) demonstrating that genotype is a poor predictor for the variation in CHC composition. A PERMANOVA approach also demonstrated a significant difference (α=0.05) between genotype in Canadian but not Norwegian females. Overall, these results support the hypothesis that CHC composition between supergene genotypes varies much more strongly in males than females. CHC profile thus appears to be a good candidate trait for female mate choice in this system, and is possibly the signal underlying disassortative mating in relation to *Cf-Inv(1)*. Intraspecific variation in pheromones is increasingly recognized in insects [28] but it rarely is connected to supergenes or chromosomal rearrangements, although a segregating putative inversion in the European corn borer moth (*Ostrinia nubilalis*) contributes to intraspecific differentiation between insect pheromone strains [29]. By tightening linkage between multiple alleles, supergene architecture may be particularly favorable for maintaining intra-specific divergence in complex traits such as chemical signaling.

For CHCs to facilitate disassortative female choice, *C. frigida* females must be able to detect male CHCs, in particular the compounds that differ between genotypes. To investigate this, we measured female perception of the different compounds using gas chromatography-electroantennographic detection (GC-EAD). During mating, males place their forelegs over the female’s head directly in contact with her antennae (Fig. 2A) so we focused on antennal perception. Reliable EAD readings could be obtained from ten Norwegian females; 2 αα, 3 αβ and 5 ββ genotypes. The females reacted to 36 compounds in a combined male extract (i.e. mixed extract from αα, αβ, and ββ males), (Fig. 2B). Conversely, male flies only sensed 3 of these compounds (based on six EAD readings, data not shown). Altogether, those results indicate that male CHCs are actively perceived by females, and provided a set of relevant candidates for sexual communication. Of the 36 female EAD active compounds, 20 could be successfully identified and integrated. These were retained for a re-analysis using individual GC-MS readings (Fig. 2B).

**Figure 2–.**
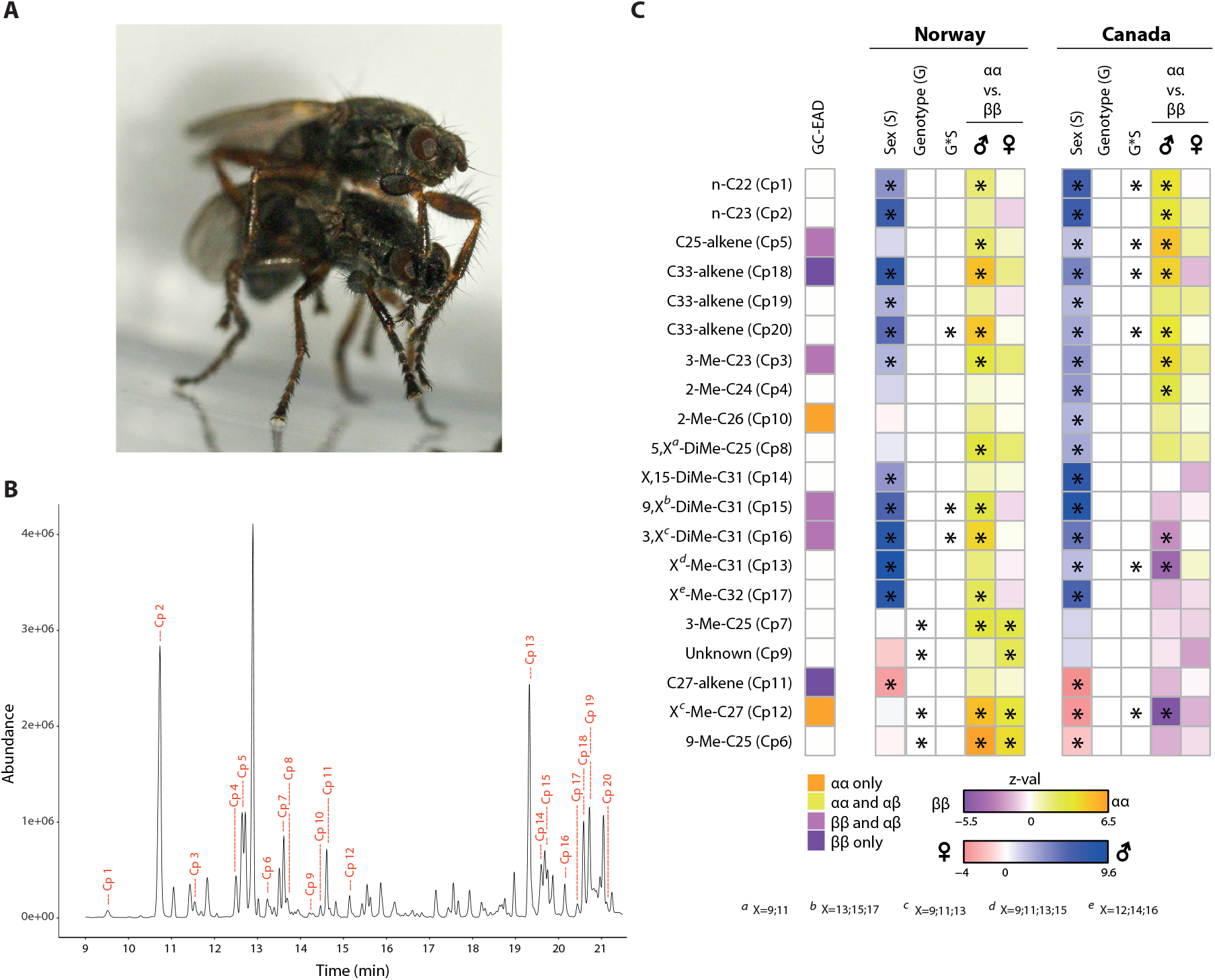
Female detection of CHCs varies by genotype. (A) Behaviour of *Coelopa frigida* during mating; note the legs of the male in direct contact with the female’s antenna. (B) Example chromatogram from an αα male with the 20 identified EAD active peaks (Cp1-Cp20) with tentative compound annotation. (C) Heatmap depicting the significance (at p=0.10 level) and strength of the effect of sex and/or genotype on the quantity different compounds. The list of compounds is restricted to the 20 compounds that are perceived by the females (as assessed by GC-EAD) and identified and integrated in CHC profiles (by GC-MS). The first column indicates compounds perceived by only a subset of genotypes.

To further understand sexual communication in relation to the supergene, we tested which compounds, among the ones perceived by females, differed between genotypes. Sixty percent of the 20 identified candidate compounds showed a consistent, significant, effect of sex between continents (Fig. 2C). Moreover, many showed an effect of genotype or an interaction between genotype and sex (Norway 7 compounds; Canada 6 compounds; Fig. 2C). The effect of genotype was mostly restricted to males (Fig. 2C) concordant with our findings on the overall CHC composition. Patterns between continents were generally consistent although 2 compounds showed significant but opposing effects (Cp12 & Cp14) and an additional 5 showed opposing effects that were significant in only one population. Intriguingly, all 7 of these compounds showed increased concentrations in αα males in Norway and ββ males in Canada. Furthermore, all 7 were identified as methyl-alkanes. Both the similar pattern and chemical similarity of these compounds indicate that there may be a simple genetic architecture to these differences. The consistency between continents indicates either parallel evolution or that the initial coupling of female perception and male traits evolved before *C. frigida* spread to other continents. However, the minor differences between populations suggest that sexual selection for male CHC composition is also ongoing independently in both North America and Europe.

If CHC composition facilitates disassortative mating with respect to *Cf-Inv(1)*, we expect that chemical perception or signal processing in females may vary between genotypes. A logistic PCA on the single female fly responses to all 36 compounds confirmed this prediction, indicating differences between the female genotypes in the odor perception (LogPCA, Fig. S2). Perception of these compounds varied between genotype and compound: four were detected exclusively by αα females, one by αα and αβ females, two others exclusively by ββ females and seven others by ββ and αβ females exclusively (Fig.2C, note that this is reduced to the 20 compounds we could identify). This suggests that *Cf-Inv(1)* controls both male traits as well as female perception of these traits. As CHC profile differ between sex but female CHCs vary little by genotype it is unlikely that a self-referencing mechanism guides mate choice relative to *Cf-Inv(1)*. Instead, disassortative mating in this system likely functions via tight linkage between male mating signals and female preferences. Although it is not possible to know how disassortative mating originally evolved, there is strong overdominance of *Cf-Inv(1)* with heterokaryotypes enjoying increased reproductive success caused by a life history tradeoff between homokaryotypes [22] as well as increased survival likely caused by masking of deleterious recessive alleles [6, 15, 16, 30]. Disassortative mating, and a linked architecture between trait and preference, are thus under strong selection, as choosing a mate with an alternate genotype ensures higher fitness. In the same vein, the pleiotropic effect of *Cf-Inv(1)*, on male traits, female preferences, and the genes underlying overdominance, results in a feed-back loop that should strengthen disassortative mating and stabilize the polymorphism as suggested by mate choice theory [3, 17] and speciation theory [31, 32].

We sought to more thoroughly explore the genetic basis of male traits and female perception using differential expression analyses. Although the pathway of CHC biosynthesis is well established [26] a recent study revealed the effect of single genes on overall CHC composition is highly complex [33]. Thus, we took a more general approach; we identified 263 candidate transcripts for both CHC biosynthesis (CHCB) and Odorant detection (OD) and tested the overall patterns of differential expression between genotypes in both sexes using previously published RNA-seq data from 17 European adults (3 αα ♀, 6 ββ ♀, 3 αα ♂, 5 ββ ♂) [34]. We performed 3 separate analyses: one with both sexes, one with males only, and one with females only. Twenty-nine transcripts putatively acting on chemical communication were significantly differentially expressed between αα and ββ, with some overlap between analyses (Table 1, Fig. S3). For the CHCB transcripts putatively involved in CHC synthesis, the signal was largely driven by males: differential expression between genotypes of 11/12 CHCB transcripts was restricted to males, and 7/12 CHCB transcripts were over-expressed in males compared to females (Table 1A). Conversely, differential expression of 11/17 OD transcripts was significant in multiple analyses and not restricted to a single sex (Table 1B).

**Table 1 -.**
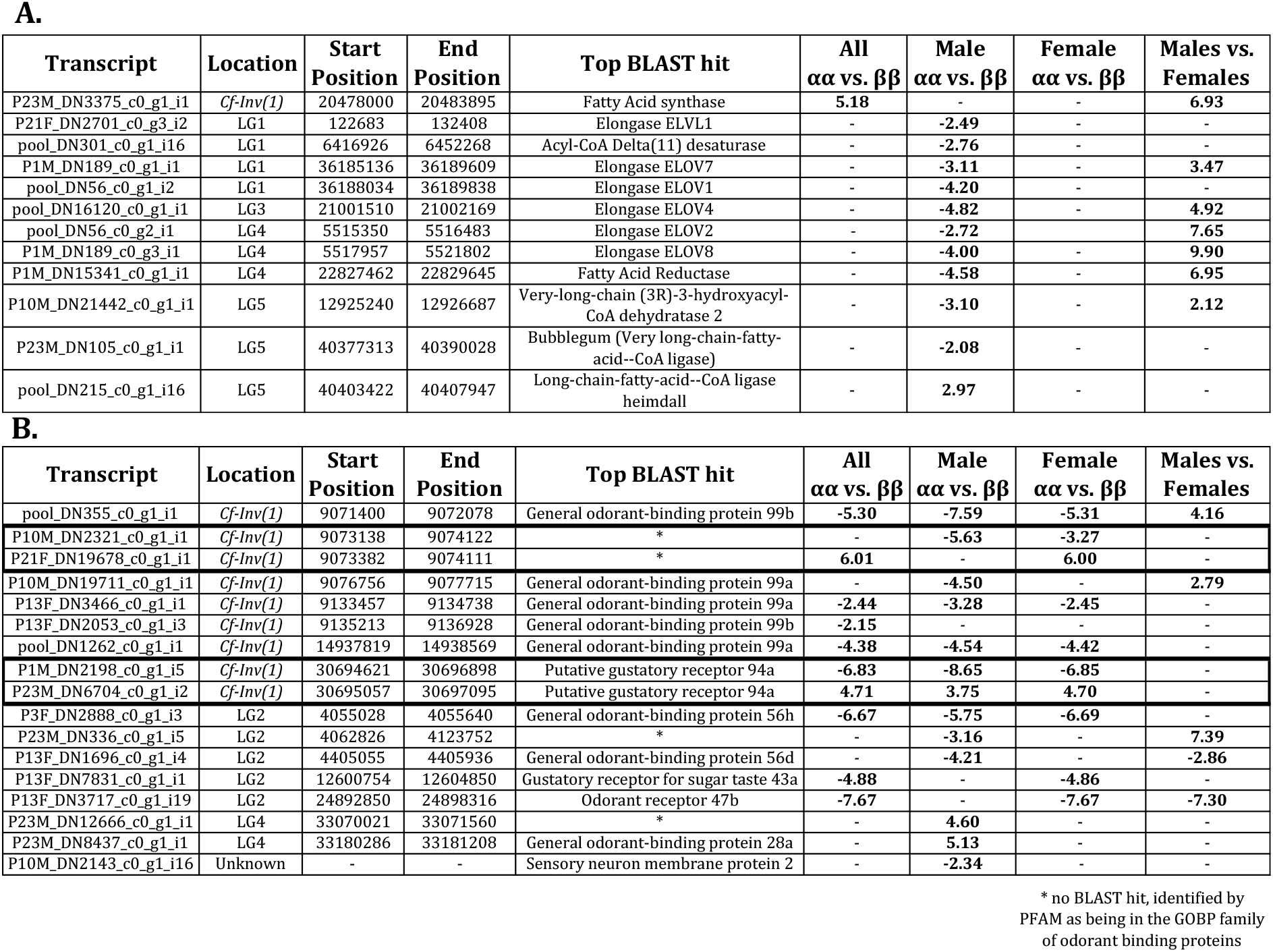
Differentially expressed transcripts involved in CHC biosynthesis (A) and odorant detection (B). Shown are the transcript id, the location in the genome, the coordinates of the transcript determined by gmap [35], the top Blast hit, and the outcomes from four differential expression analyses. Only log2fold values (bolded numbers) for significantly differentially expressed transcripts are shown. Black boxes indicate pairs of duplicated and divergent transcripts.

To ask whether the differentially-expressed genes were located within *Cf-Inv(1)*, we mapped our candidate transcripts to version 1.0 of the *C. frigida* genome [36]. The two groups of genes showed strongly different patterns of localization. CHCB differentially-expressed transcripts were widespread in the genome, with only one transcript mapping to the supergene, while nine OD transcripts mapped to *Cf-Inv(1)*. This is significantly more than expected (52.9% of differentially-expressed OD transcripts compared to 17% of all tested transcripts; Fisher’s Exact Test p=0.0053). Therefore, on the signal side, although the effect of *Cf-Inv(1)* on CHC composition was, as predicted, limited to males for both the phenotype and gene expression, it appears to be *trans* in relation to *Cf-Inv(1)* itself. However, we still found clustering of CHCB loci, similar to what has been found in *Heliconius* butterflies [37]. As males and females share a genome, sex specific changes in expression via cascading effects are more likely [38]. In contrast, on the reception side, the excess of OD genes within *Cf-Inv(1)* indicates a disproportionate effect of *Cf-Inv(1)* on chemical perception.

Next, we examined the distribution of odorant detection genes across the genome regardless of expression status. We further subset our putative odorant detection genes to look at odorant receptors (ORs), gustatory receptors (GRs), and odorant binding proteins (OBPs) as these were found to be prime candidates for mate choice in *Heliconius* [39]. We found 63 transcripts that mapped to the genome (Table S3). The distribution of these transcripts was non-random, as several transcripts with similar or identical annotations were clustered at close proximity in blocks of 2 to 10 transcripts (Fig. 3A, Table S3). Such a clustering may be the result of tandem duplications, and is frequently observed for odorant-binding proteins and chemosensory receptors in insects [40–43]. Evolution by tandem duplication and subsequent divergence (e.g., the birth death model of multi-gene families [44]) is proposed to have generated the large and diverse OD gene families identified in insects [41, 43, 45, 46]. While there was not an excess of OD genes within *Cf-Inv(1)* compared to its size, it had an excess of paired transcripts with overlapping coordinates (labeled with A/B Fig. 3A).

**Figure 3–.**
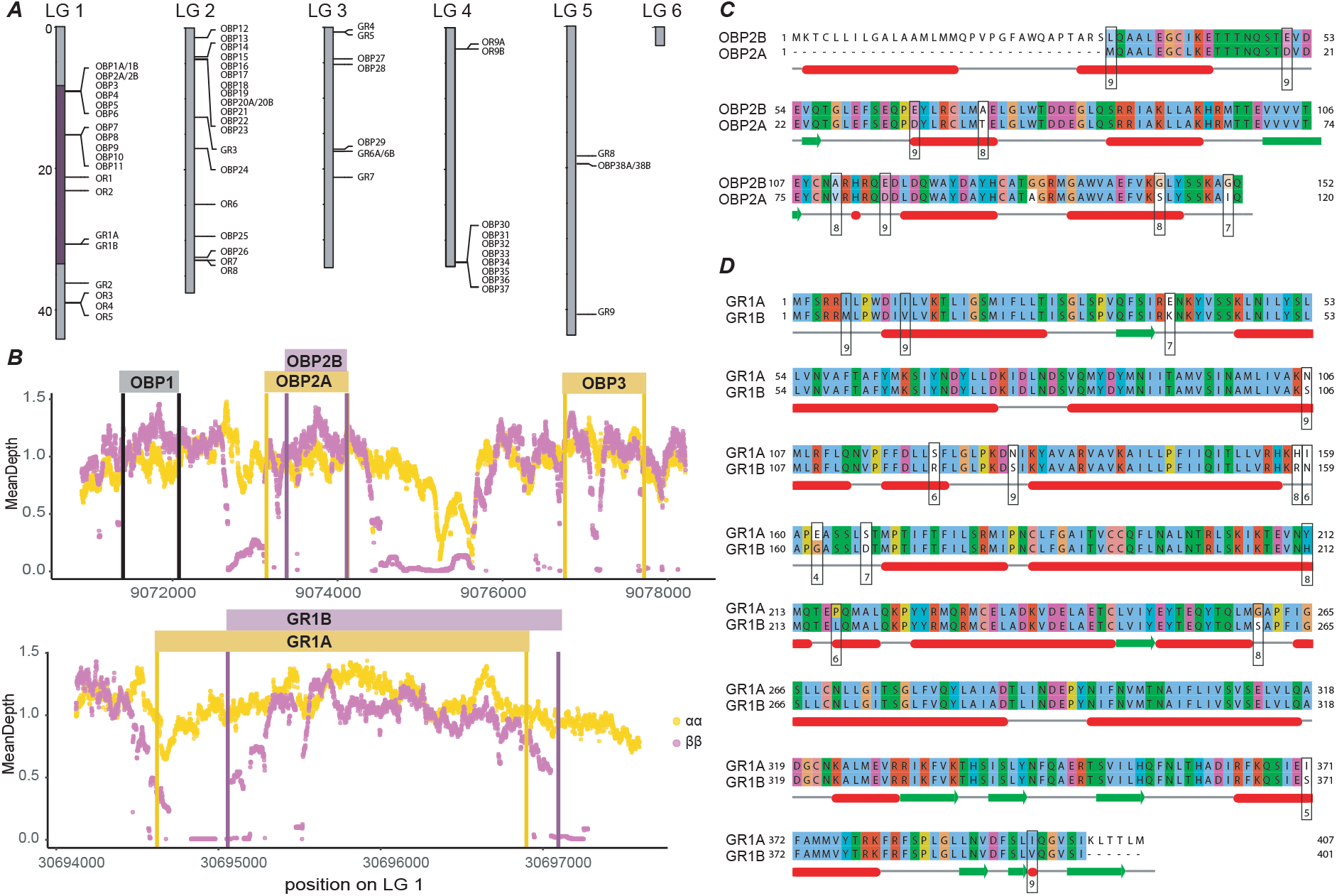
Divergence of odorant detection genes within the supergene. (A) Clusters of odorant detection genes identified in the *C. frigida* genome. Putative genes were labeled as OR - odorant receptor, GR - Gustatory receptor, OBP - odorant binding protein. Numbers correspond to their order in the genome starting from LG1 position 1. Overlapping pairs were labeled with the same number and title but with A and B afterwards. Visualization was done with karyoploteR [47] (B) Coverage: Average depth of sequencing across genotypes at all positions along the focal genomic regions. Expected depth is around 1.1X. Bars indicate the position of the transcripts. (C,D) Protein alignment of OBP2A/B (C) and GR1A/B (D). Amino acids are colored to show biochemical properties using the Clustal × coloring scheme. Protein structure predicted by Jpred is shown below with red bars indicating helices and green arrows indicating sheets. Below that is the AMAS conservation score with higher values indicating more conserved amino acids [48].

We found 6 pairs of transcripts with overlapping coordinates, 3 of which mapped within *Cf-Inv(1)*. This pattern could be due to isoforms, exon duplications, errors in transcriptome assembly, or divergence. Two of these pairs (9.07Mb and 30.69Mb) were of particular interest as they showed opposite expression patterns (Table 1). One transcript of each pair was assembled from an αα individual and the other transcript from a ββ individual, indicating that they probably represent alternative alleles of the same ancestral genes. Reusing previously-published whole-genomes sequences [36], we observed that both genomic regions are characterized by high genetic differentiation between α and β (F_ST_=0.96-0.97 and d_xy_ = 0.04-0.07 compared to F_ST_=0.87 and d_xy_=0.02 in *Cf-Inv(1)* overall), and heterogeneous coverage in ββ (Fig. 3B), which may reflect either low mapping of those sequences on a αα reference genome, or ββ-restricted deletions. Comparing the protein sequences of these pairs showed striking divergence in both amino acid identity and biochemical properties (Fig.3C, D). The third pair within *Cf-Inv(1)* was also somewhat divergent, but less so (Fig. S4). This indicates that the overlapping pairs within *Cf-Inv(1*) are due to divergence between supergene haplotypes. We note that, after merging these pairs there was still a marginally significant excess of differentially expressed OD transcripts within *Cf-Inv(1)* (Fisher’s Exact Test p=0.0996). Thus, we found divergence in both coding sequences and expression. In most insects diversification takes place between species or tandem duplicates [41, 43], however, in *Coelopa frigida*, we propose that divergence may have taken place within species, between arrangements of *Cf-Inv(1)*, although data from sister species is necessary to better understand the evolutionary history of those genes. Overall, the duplication and divergence of OD genes between the two arrangements of the supergene is comparable to patterns observed between sister-species of insects [49]. This is in line with fast evolution of odorant-binding proteins and chemosensory genes [43] and suggests that supergenes provide a genetic architecture favoring the rapid divergent evolution of mate choice including at the intra-specific level.

## Conclusion

Overall, our findings highlight the importance of genetic architecture on the evolution of intra-specific diversity and the persistence of supergene-associated polymorphism. At a coarse scale, the linkage between hundreds of loci within the supergene is putatively responsible for the selective pressure that favors disassortative mating by underlying heterozygote advantage while the linkage between the genes underlying male signal and the genes underlying female perception likely provides an ideal architecture for the persistence of disassortative mating itself. Moreover, at a finer scale, the duplication of odorant detection genes within the supergene region possibly contributed to the divergence of signal reception and the rapid evolution of mate preference. Reciprocally, disassortative mating coupled with other mechanisms of balancing selection, such as spatially varying selection, preserve the coexistence of supergene haplotype across long time scales, further enhancing the accumulation of divergence.

## Methods

### Collection and rearing of flies for CHC analysis

Wild *C. frigida* adults were collected in September 2018 at Kamouraska, QC, Canada (47.56294, −69.87375) and transported to Laval University. Wild *C. frigida* larvae were collected from Østhassel, Norway (58.07068, 6.64346) in March 2018 and transported to Tjärnö Marine Laboratory, Gothenburg University in Strömstad, Sweden. Laboratory conditions were standardized between Sweden and Canada and all files were raised in a temperature controlled room at 25°C with a 12-hr/12-hr light–dark cycle. The Canadian adults were allowed to lay eggs on standardized lab wrack (approximately 50% Fucoids and 50% Kelp) and the subsequent generation was used for CHC analysis (*i.e*. the focal generation). The Norwegian larvae were allowed to mature in their natural wrack and the emerging adults were used to generate the focal generation, which was reared on standardized lab wrack (approximately 50% Fucoids and 50% Kelp).

For the focal generations, as larvae pupated, they were transferred to individual 2 mL tubes with a small amount of cotton soaked in a solution of 0.5% mannitol to provide moisture and food for the eclosing adult. This ensured virginity in all eclosing flies. The tubes were checked every day, and the date of eclosure of each fly was noted. Flies mature approximately 24 hr after eclosion at 25 degrees so, two days after eclosure, we froze them at −80°C. To genotype each adult at the inversion, we extracted genomic DNA from one foreleg and performed a diagnostic SNP assay involving a PCR step amplifying the *Adh* gene, and a digestion step with two restriction enzymes targeting SNPs fixed between arrangements (described in Mérot et al 2018). Frozen flies from Canada were sent on dry ice to Tjärnö Marine Laboratory for CHC extraction.

### CHC Analysis

Frozen flies were allowed to defrost and dry for 10 minutes, as moisture can affect lipid dissolution. Each fly was then placed in a 1.5 ml high recovery vial containing 300 µl of *n*-hexane, vortexed at a low speed for 5 seconds, and extracted for 5 minutes. Afterwards flies were removed from the vial and allowed to air dry before they were weighed on a Sartorius Quintix 124-1S microbalance to the nearest 0.0001 grams. Extracts were evaporated until dryness under a stream of nitrogen and stored at −20° until GC-MS analysis. Before analysis, extracts were redissolved in 20 µl of *n*-hexane containing 1 µg/ml *n*-nonane (Sigma-Aldrich) as an internal standard and vortexed at maximum speed for 10 seconds. Extracts of 25 flies per population (Norway and Canada), sex and karyotype were analyzed on a GC (Agilent GC 6890) coupled to a MS (Agilent 5973 MSD) using a HP-5MS capillary column (Agilent, 30m × 0.25mm ID, 0.25µm film thickness). Two microliters were injected in a pulsed splitless mode. Inlet and MSD transfer line temperature was set isothermal to 280°C. The oven temperature was programmed to 190°C for 3 min, increased to 280°C at a rate of 8°C/min, then increased to 325°C at a rate 4°C/min and finally held at this temperature for 14.5 min. Helium was used as the carrier gas at a constant flow of 0.5 ml/min. The mass spectrometer was operated in scan mode scanning ions from 40–700 amu (2.24 scans s^−1^) with an electron ionization at 70 eV, a source temperature at 230°C and quadrupole temperature at 150°C and an initial 3 min solvent delay.

The total ion chromatograms were quality checked, de-noised and background subtracted using CODA and Savitzky-Golay smoothed before peak detection and peak area integration in OpenChrom®. Peaks were aligned using the R-package ‘GCalignR’[50]. Peaks with more than 80% missing values and max peak areas below 1 × 10^6^ were excluded from the data set. The peaks kept in the final peak list were then manually revised and missing peaks were manually integrated when applicable or imputed by a random forest algorithm using the R-package “missforest” [51]. Unfortunately, the chromatograms revealed co-eluting contaminants from cap material for the majority of the Norwegian samples. Different caps were thus used for the remaining samples. The contaminated Norwegian samples were excluded from global CHC analysis (PCA, OPLS-DA) but included for analysis targeting homologous peaks identified in all samples.

#### Statistical analyses

For the analysis of CHC profiles, peak areas were normalized on the internal standard peak area and the weight of the fly. For the multivariate analysis, all data were additionally mean centered and unit variance scaled. Extreme outliers were identified by deviations in the orthogonal and score distance, and they were removed prior to analysis. Balanced sample sets were visualized with PCAs for Canadian females (αα /ββ=22), Canadian males (αα /ββ=22), Norwegian females (αα /ββ=10) and Norwegian males (αα /ββ=5). Differences between αα and ββ karyotypes were assessed by OPLS-DA using the “ropls”-package in R [52] and PERMANOVA on Euclidean distances using the “vegan” package in R [53]. OPLS-DA model significance was estimated by permutation (number of iterations=999). Candidate compounds for differentiation between karyotypes were determined from variables of importance for OPLS-DA projection (VIP scores) [54] combined with an univariate analysis using a false discovery adjusted significance level of α = 0.05 following the Benjamini and Hochberg’s FDR-controlling procedure (Benjamini & Hochberg, 1995). Compounds with VIP>1 and adjusted p-value<0.05 were tentatively annotated by their mass spectra and retention time index based on a C21-C40 *n-*alkane standard solution (Sigma-Aldrich).

### Gas chromatography and electroantennographic detection (GC-EAD)

All GC-EAD analyses were performed at the Nature Research Centre in Vilnius, Lithuania. Two µl of *C. frigida* male CHC extract was injected splitless into a gas chromatograph (GC) Clarus 500 (Perkin Elmer, Shelton, CT, USA) equipped with an Elite-5 column (30 m × 0.25 mm× 0.25 µm, Perkin Elmer, Shelton, CT, USA). The effluent from the column was split into two equal parts, one part was transferred to flame ionization detector (FID) and the other part to electroantennographic detector (EAD). Two signals from an olfactory active compound were registered simultaneously, the first one recorded by the GC detector FID and the second one recorded from a fly antenna mounted to EAD. The injector, FID, and EAD transfer line temperatures were set isothermal at 280°C, 280°C and 240°C, respectively. EAD transfer line temperatures were controlled by a temperature controller, TC-02, (Syntech, Buchenbach GmbH, Germany). The same oven temperature program was used as in the GC-MS analysis (see above). Hydrogen was used as the carrier gas (1 ml/min) and the makeup gas (5 ml/min). EAG registration was carried out using a signal connection interface (IDAC 4, Ockenfels Syntech GmbH, Buchenbach, Germany). Data storage and analysis were carried out with a PC-based interface and software package GcEad V. 4.4 (Synthech, Ockenfels Syntech GmbH, Buchenbach, Germany).

The flies used in GC-EAD analyses were not chilled prior to testing. The head of an insect was removed with scissors, and the tip of the left antenna arista was cut off. A glass capillary indifferent electrode filled with 0.9% NaCl saline (Ilsanta, Vilnius, Lithuania) and grounded via a silver wire was inserted into the hemocoel of the cranial cavity, and the recording electrode was connected to the cut tip of the arista. Activated charcoal filtered and humidified constant airflow at a rate of 1 L/min passed over the antenna through a glass tube (ø 7 mm) positioned 0.5 cm from the antenna. Only unmated adults (males and females) from the Norwegian population were used. Adults were kept allowed to feed on seaweed for 24 h before the testing. The test was carried out at 22 ± 2°C. Two or more successful GC-EAD recordings were obtained for consideration of electrophysiologically active CHCs. As GC-EAD recordings were binary data we tested for an effect of genotype with a logistic PCA using the logisticPCA package [55] implemented in R.

#### Analysis of EAD active compounds

Retention indexes were calculated from the EAD/FID chromatograms and translated into expected retention times in the GC-MS chromatograms. By comparison of the expected retention times together with the GC-FID elution patterns, the EAD active peaks were identified in the GC-MS chromatograms. We then focused on this subset of EAD active compounds and extracted their peak areas manually from the GC-MS chromatograms for the αα and ββ karyotypes. αβ karyotypes were excluded for this analysis since the multivariate analysis showed that the CHC profiles of AB are intermediates between αα and ββ. The Norwegian samples with contaminants from cap material could be included in this analysis after verification that the retention times of the EAD active compounds differ from the retention times of the observed contaminants introduced by the cap material. The EAD active compounds were normalized on the internal standard peak area and the weight of the fly prior to analysis. For each EAD active compound, differences between karyotypes within sex and differences between males and females were analyzed using a glm approach implemented in R with sex, genotype, and their interaction as potential factors. Contrast statements to specifically test for the difference between αα and ββ in males and females were implemented post-hoc.

### Differential expression analysis

We used several methods to identify candidate genes for CHC biosynthesis and odorant detectors. First we searched flybase (flybase.org) in December 2019 for the gene groups “odorant binding proteins” (52 hits) and “chemoreceptors” (121 hits). At the same time we searched the uniprot database for the term “Cuticular hydrocarbon biosynthetic process” (85 hits). All isoforms from matches were downloaded as protein fasta files. We also added 17 candidate genes from a recent GWAS study in *Drosophila melanogaster* that had been functionally validated (Dembeck et al 2015, elife). We used tblastn with a e-value of 1e-3 and a maximum of 5 target sequences to find orthologs of all of these candidate genes these in our transcriptome. The *C. frigida* transcriptome has previously been annotated and so we also searched for contigs annotated with at least one of the GO terms: “sensory perception of smell”, “sensory perception of chemical stimulus”, “cuticle hydrocarbon biosynthetic process”, “fatty acid biosynthetic process”, “odorant binding”, “response to odorant”, and “olfactory receptor activity”.

We used previously reported data to examine differential expression at candidate genes for CHC biosynthesis as well as odorant detection. All details on the generated data set can be found in Berdan et al. 2021 [34]. This data set contains *C. frigida* from a several Swedish and Norwegian populations. We subset this data set to only retain data from adults (3 αα males, 3 αα females, 5 ββ males, and 6 ββ females). We used previously generated count matrices and used DESeq2 [55] to determine differentially expressed genes between αα and ββ in males, females, and a combined analysis with both sexes. After the DESeq2 analysis but before applying an FDR correction the results were subset to only include our candidate genes. Conventional thresholds (log2 fold change > 2, adjusted p-value (FDR) < 5%) were used to identify differentially expressed transcripts within our candidate subset.

### Analyses of clusters of OD transcripts

For examination of genomic locations, we subset our list to only include transcripts that had a blast match containing one of the following terms ‘odorant binding protein’, ‘gustatory receptor’, or ‘odorant receptor’ or a pfam annotation of GOBP (PF01395). Proteins from the two pairs within *Cf-Inv(1)* (Fig. 3B) were predicted using transdecoder (http://transdecoder.github.io) implemented in Trinity [56] using the longest ORF option. Complete proteins were aligned using the EMBOSS Needle global aligner run on the EMBL-EBI website (https://www.ebi.ac.uk/Tools/psa/emboss_needle/) and visualized using Jalview [57]. Conservation between sequences was calculated using the AMAS method [48] and protein structure was predicted using Jpred4 (http://www.compbio.dundee.ac.uk/jpred/) [58].

Coverage, F_ST_ and dxy between genotypes were extracted from previously-published analyses [36], sub-setting the dataset to 144 αα and 144 ββ wild-caught *Coelopa frigida*. Briefly, low-coverage whole genome sequences were aligned to the reference genome (assembled from a αα individual), and coverage, polymorphism and differentiation were analyzed in ANGSD.

## Acknowledgements

We thank A. Kinnby, E. Riedel, G. Cervin, and G. Nylund for help with fly rearing and seaweed acquisition and T. Flatt for helpful comments that improved the manuscript. C.M. and L.B. would like to thank F. Larochelle for establishing the conditions for rearing the flies and S. Bernatchez and M. Leitwein for helping in the field. E.L.B. was supported by a Marie Skłodowska-Curie fellowship 704920 – ADAPTIVE INVERSIONS and gratefully acknowledges funding from Helge Ax:son Johnsons Stiftelse. C.M. was supported by fellowship from Fonds de Recherche du Québec (FRQS260724, FRQNT200125) and a Banting postdoctoral fellowship (#162647). Research on the Canadian flies was funded by a Discovery research grant from the Natural Sciences and Engineering Research Council of Canada (NSERC) to L.B. H.P. gratefully acknowledges funding from The Swedish Foundation for Strategic Environmental Research MISTRA (grant no. 2013/75) from the Swedish Foundation for Strategic Research (Sweaweed). Electrophysiological studies were funded by a Lithuanian state grant to the Nature Research Centre within Research program III, Molecular, genetic and evolution mechanisms of functioning and adaptation of biological systems. The gene expression analysis was enabled by resources provided by the Swedish National Infrastructure for Computing (SNIC) partially funded by the Swedish Research Council through grant agreement no. 2018-05973.

