## Supplementary Figures for "A supergene in seaweed flies modulates male traits and female perception"

### Supplemental Figures

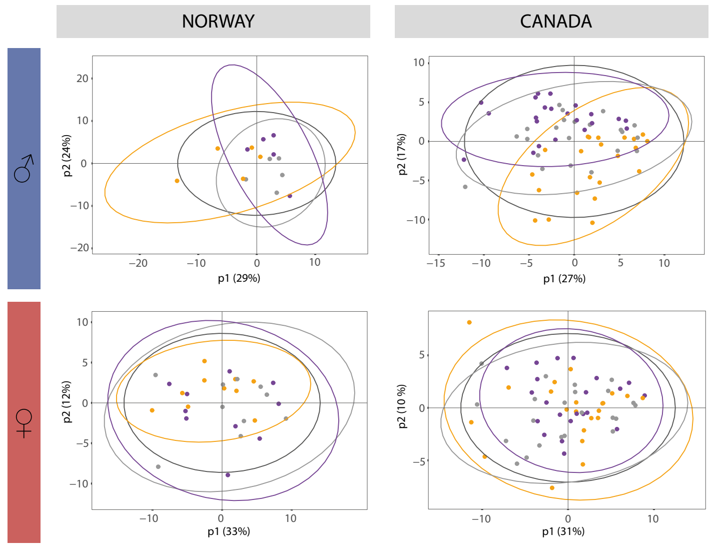


**Figure S1** - Principal Component Analysis of CHC composition in Norwegian and Canadian populations. Figures are divided by population and sex and colored by genotype: orange - αα, grey - αβ, purple - ββ.

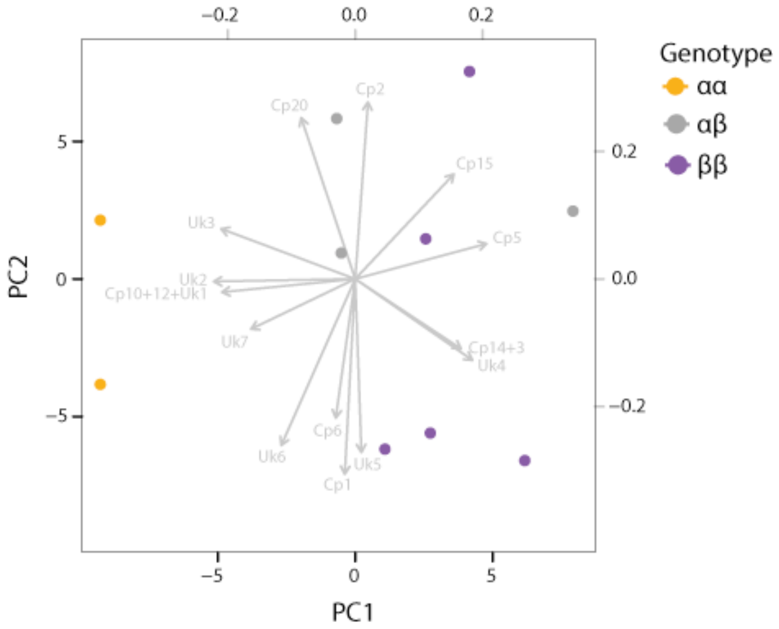


**Figure S2** - LogPCA of binary EAD responses of females. Only compounds showing an absolute correlation => 0.15 with either component 1 or 2 are shown. Samples are colored by genotype: orange - αα, grey - αβ, purple - ββ.

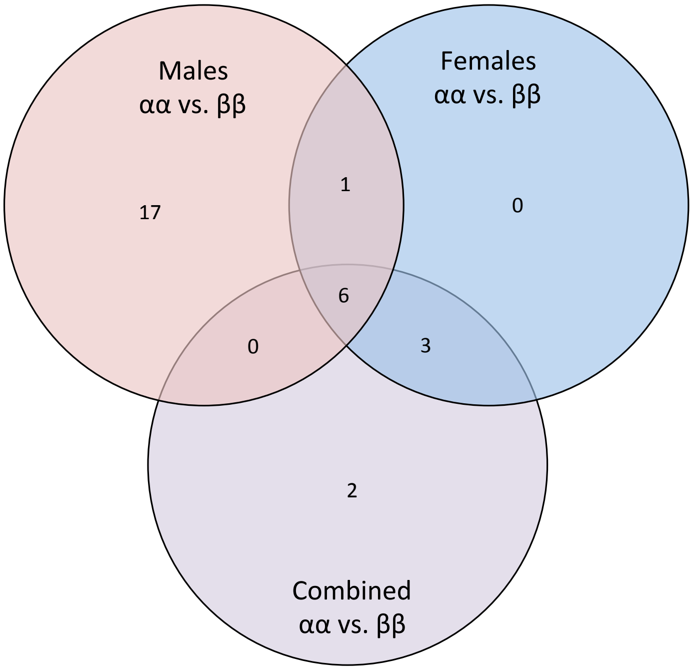


**Figure S3** - Overlap between differential expression analyses. Shown is the overlap of transcripts that are significantly differentially expressed between αα and ββ in at least one analysis. Circles are colored by analysis: red - females, blue - males, purple-combined.

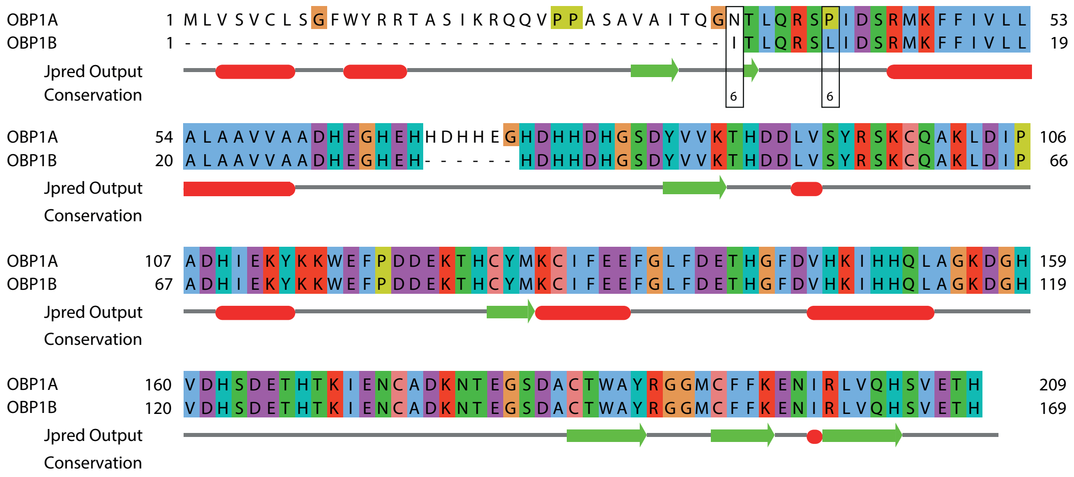


**Figure S4** - Alignment of protein sequences OBP1A and OBP1B. Amino acids are colored to show biochemical properties using the Clustal X coloring scheme. Protein structure predicted by Jpred is shown below with red bars indicating helices and green arrows indicating sheets. Below that is the AMAS conservation score with higher values indicating more conserved amino acids [55]. Note that OBP1B is only a 5’ partial protein.
